# Role of hemocytes in the regeneration of germline stem cells in *Drosophila*

**DOI:** 10.1101/2020.08.31.275255

**Authors:** Virginia Beatrix Varga, Fanni Szikszai, Janka Szinyákovics, Anna Manzéger, Gina Puska, Tibor Kovács, Tibor Vellai

## Abstract

Cellular regeneration, which relies on extensive restructuring of cytoplasmic materials, is an essential process to restore tissues and organs lost during aging, degenerative diseases and injury. At early stages of *Drosophila* spermatogenesis, when cellular constituents are intensely remodeled, there are two different populations of stem cells, the somatic stem cells and the germline stem cells (GSCs). GSCs divide by asymmetric division to give rise two distinct daughter cells. One of them will leave the stem cells’ niche and differentiate into spermatogonial cells (SCs). Both aging and cellular stress can lead to the loss of GSCs. Lost GSCs can be restored by dedifferentiation of SCs into functional GSCs. In other tissues, macrophages provide specific conditions for cellular transformation. Here we examined the potential role of immune surveillance cells called hemocytes during dedifferentiation of SGs into GSCs. We found an elevated number of hemocytes during this dedifferentiation process. Immune depletion of hemocytes decreased the regeneration capacity of germline. We also show that autophagy, which plays a pivotal role in cellular differentiation by eliminating unwanted, superfluous parts of the cytoplasm, becomes upregulated in dedifferentiating SCs upon JAK-STAT signaling emitted by hemocytes. Furthermore, these immune cells regulate expression of Omi/HtrA2, a key regulator of apoptosis in early spermatogenesis. Together, we suggest that hemocytes have important functions in the dedifferentiation process of GSCs.

## Introduction

Regeneration is a cellular process by which cells lost during aging, injury and various cellular insults can be effectively restored to maintain tissue functioning. The regeneration process is based on cell fate transformation mediated by remodeling of cellular constituents^1^. Cellular regeneration can be achieved by three distinct major mechanisms: 1) dedifferentiation of a terminally differentiated somatic cell into an unspecialized blastemal cell having the capacity to further proliferate and the daughter cells differentiate into the lost terminally differentiated cell type, 2) direct transdifferentiation of a specialized somatic cell into another specialized cell type, and 3) reprogramming from local somatic stem cells. During dedifferentiation, the affected specialized somatic cell regains its potency to differentiate into a cell type that characterizes the original lineage when differentiating into the blastema^2^. A tractable tissue model for studying dedifferentiation is represented by the zebrafish caudal fin, which can effectively be regenerated followed by amputation within a week. At the site of amputation, unipotent progenitor cells are generated which constitute the blastema (a mesenchymal cell mass serving the source of the regenerating tissue)^3^.

According to the literature, macrophages, which are specific immune cells, play an important role in tissue regeneration mediated by dedifferentiation. For example, these cells participate in the regeneration process of zebrafish caudal fin and amphibian limb^4,5^. It was also shown that residual macrophages affect the proliferation of somatic stem cells. For further studying the role of macrophages in tissue regeneration by dedifferentiation, the *Drosophila* testis may also serve as an ideal *in vivo* model system, since the Drosophila melanogaster male germline stem cell niche is one of the best characterised stem cell system^6,7^. In the fruit fly, specific immune cells called hemocytes involve plasmatocytes, lamellatocytes and crystal cells^8^. Plasmatocytes are the dominating hematocyte population during all developmental stages, and capable of phagocyting. Regarding their function, these cells are most similar to mammalian macrophages. Plasmocytes, together with crystal cells, generate the von Willebrand factor, also called hemolectin (Hml), which we used here as a marker for detecting these cells in the regeneration process^9^. Hemocytes also function to eliminate apoptotic bodies of spermatogonial origin through phagocytosis^10^. Plasmocytes are able to emit a ligand, Upd3 protein, which serves as a ligand for the JAK-STAT signaling pathway^11^. Hence, hemocytes play a role not only in humoral immune response but also in processes associated with JAK-STAT signaling^12^. It has been demonstrated that during proliferation of *Drosophila* intestinal stem cells (ISCs) hemocytes produce Upd ligands and Dpp (Decapantaplegic), thereby affecting both JAK-STAT and BMP (bone morphogenetic factor) signaling^13^. According to results provided by Van de Bor and colleagues, hemocytes participate in shaping and maintaining the microenvironment of female GSCs, and regulating their number and differentiation by producing signaling factors and cell adhesion molecules^14^.

The number of GSCs can be lowered in response to certain insults such as stress and aging. The lost cells can be restored by dedifferentiation of SGs under favorable conditions^15^. In the apical part of the *Drosophila* testis, Hub serves as a regulatory center for the stem cell niche that consists of somatic origin, postmitotic cells. Hub binds to GSCs and somatic cyst stem cells (CySCs) through cell adhesion molecules, and the coordinated operation of the three cell populations is required for the continuous production of sperms. In this region, Hub produces Upd ligand for JAK-STAT signaling, hence primordial germ cells found only at the apical part of the gonad can adapt GSC fate, and bind directly to Hub^16^.

We performed experiments on the *Drosophila* testis, which is an ideal model for studying the regeneration process via dedifferentiation because GSC regeneration can be activated under laboratory conditions. In a transgenic fly strain. *bam* driven by a heat inducible promoter ectopically expresses the main germline regeneration factor Bam (bag of marbles)^17^. Bam is thus expressed in GSCs and induces their leaving the germ line niche and differentiation into SGs. By finishing ectopic expression of Bam, dedifferentiation of SGs into GSCs can be activated. These newly produced GSCs then initiate to divide asymmetrically and generate daughter cells that eventually create sperms. This process is termed as the regeneration of early spermatogenesis^18^. To assay the process, we applied an immunohistochemical method that indicates hub cells by red (anti-FasciclinIII) and germline cells by green (anti-Vasa) signals (Fig. 1). GSCs bind to hub by cell adhesion molecules which are indicated by white dotted lines. Conditions are as follows: control (maintained at 25°C), *bam* overexpressing, and 6 and 24 hour-regeneration.

**Figure 1.**
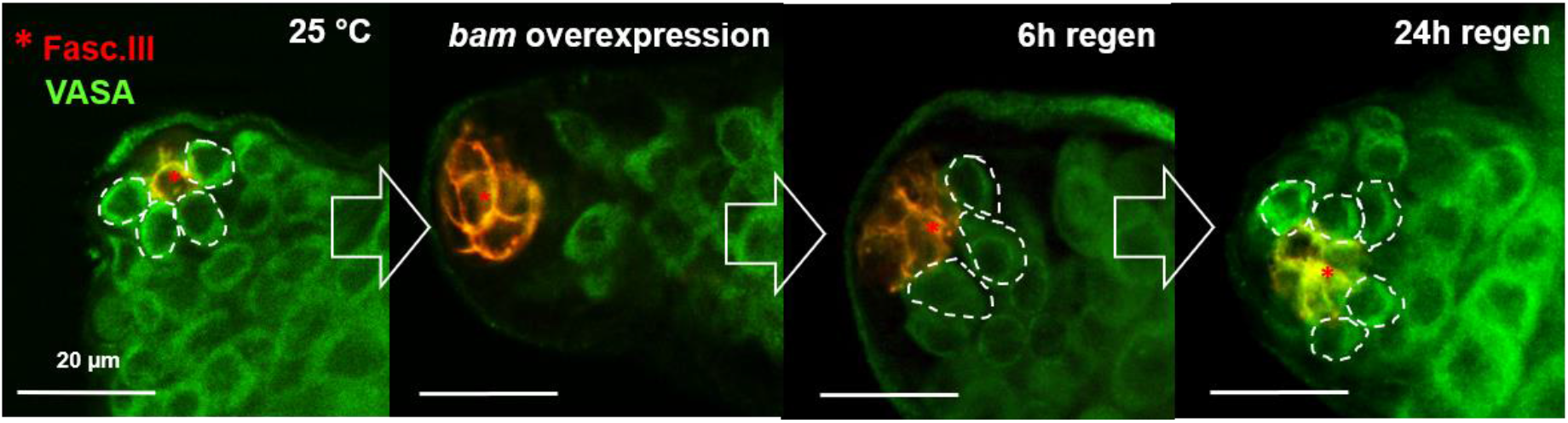
Dedifferenetiation of GSCs in the Drosophila testis. The process is based on hs-bam, bamGal4 expression system,in which Bam serves as the germline differentiation factor. Under normal condition Bam is blocked in the GSCs, while using our expression system ectopic expression promotes differentiation in the germline. Termination of ectopic expression makes regeneration of the germ stem cells possible. We examined regeneration after 6 hours and 24 hours after termination. Dotted lines indicate GSCs. Anti-FasciclinII (red) shows hub, anti-VASA (green) indicates germline cells.

## Objectives

Our aim was to uncover novel cellular and molecular factors required for GSC regeneration in the *Drosophila* testis. We also examined whether hemocytes have a role in this process because our preliminary results have indicated that inhibiting hemocyte functions decreasess the number of GSCs. We monitored a mitochondrial serine-protease involved in cell death during spermatogenesis, Omi/HtrA^19^

## Results

We previously examined lysosomal degradation during germline regeneration in the *Drosophila* testis. These experiments revealed large, multipodial cellular structures that are characteristics of hemocytes (Fig. 2).

**Figure 2.**
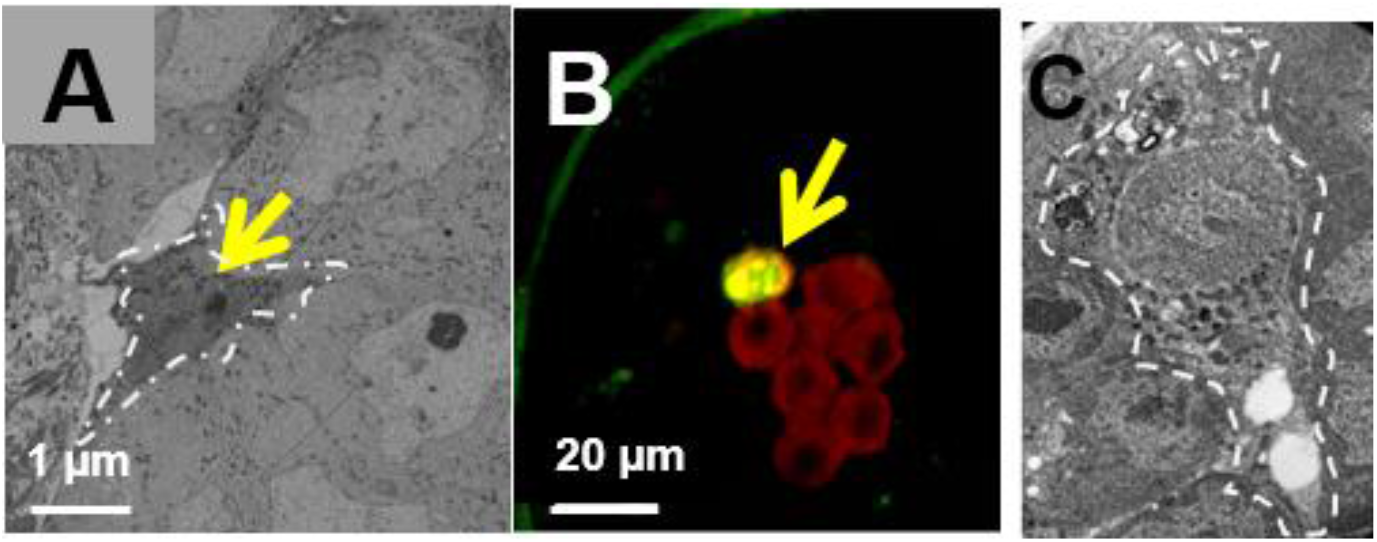
*Hemocytes are found in the* Drosophila *testis*. **A**, **C**, Specific podial structures of hemocytes are visible. These structure are capable of surrounding debris of apoptotic spermatogonial cells and their phagocytosis. **B**, Hemolectin (Hml-GFP) labels hemocytes (a fluorescent image). Hemocytes indicated by dotted lines and yellow arrows are recruited during the regeneration process and can phagocyte SGs.

It was previously described that residual macrophages can be found in the *Drosophila* testis and these are identified as hemocytes^20^.We confirmed the presence of these immune cells in the testis by using hemocyte-specific cell adhesion markers such as hemolectin (Hml)^21^. Applying a Hml-GFP transgene, we observed that hemocytes are recruited to the site GSC regeneration (Fig. S1). We also wondered whether inhibiting hemocyte function affects the regeneration capacity of GSCs. For that, germline cells were indicated by anti-Vasa (green), while hub cells were labeled by anti-FasciclinIII (red) staining (Fig. 2). Elimination of immune cells was established by overexpressing a proapoptotic gene, *reaper* (rpr)^22^. According to our results, in the absence of hemocyte function GSC regeneration becomes compromised (Fig. 3).

**Figure 3.**
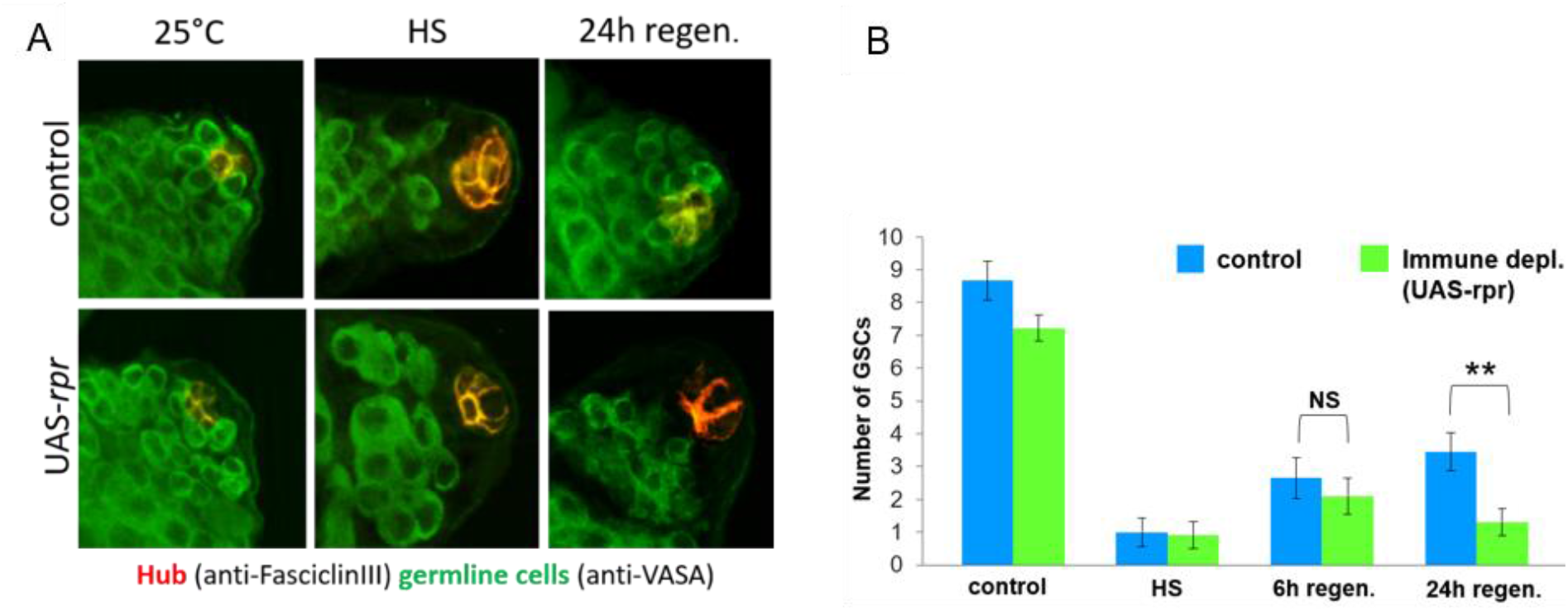
Hemocytes participate in GSC regeneration. **A**, Control *bam* overexpression (*HS*) and 24-hour regeneration in control vs. repair overexpression (*UAS-rpr*) genetic backgrounds. Both *HS* and *UAS-rpr* lead to defects in the regeneration process. GSCs indicated by dotted lines directly bind to hub. Anti-Vasa (green) staining indicates germline cells, anti-FasciclinIII (red) staining shows hub cells. **B**, Statistical analysis of immune histochemical results. Control vs. immune-depleted animals were compared. A significant difference (**: P<0.001) is seen at 24 h regeneration. NS: non-significant. ±S.D. indicates standard deviation.

It was shown that hemocytes regulates processes via JAK-STAT signaling^23^. This prompted us to analyze their potential function in GSC regeneration. The expression of *chinmo* serving as a target gene for JAK-STAT signaling was examined by RT-qPCR^24^. Relative mRNA levels were found to gradually increase during the regeneration process (Fig. 4).

**Figure 4.**
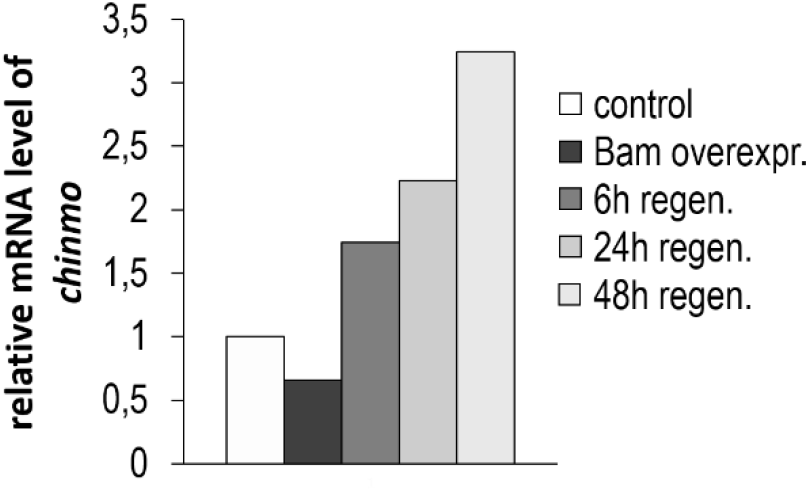
*The expression of* chinmo, *a target gene for JAK-STAT signaling, progressively increases during GSC regeneration*. During regeneration we examined increased expression level of chinmo (JAK/STAT target gene) by qPCR.

According to our results JAK-STAT signaling is active during GSC regeneration. We next asked whether hemocytes contribute to the activation of the signaling pathway in this process. *Upd1* and *Udp3* genes encode ligands for JAK-STAT signaling and were downregulated by RNA interference specifically in immune cells using HmlGal4 (hemocyte-specific) driver. Using immunohistochemistry, we showed that Upd1 and Upd3 deficiency interferes with the regeneration capacity of GSCs (Fig. 5).

**Figure 5.**
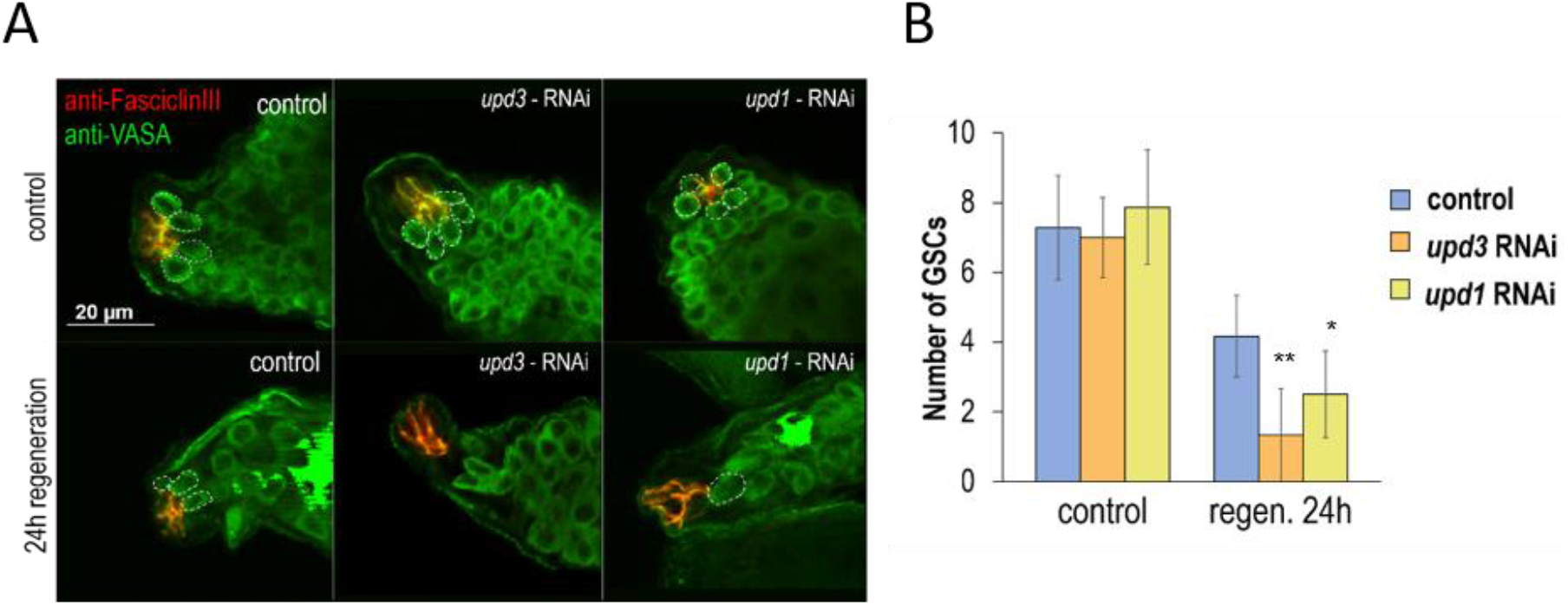
Dedifferenetiation of SGs into GSCs relies on JAK-STAT signaling emitted by hemocytes. ***A**, Anti-FasciclinIII staining (red) indicates hub cells, while anti-Vasa stining (green) labels germline cells. SG dedifferentiation is lowered in genetic backgrounds defective for Upd1 and Upd3 (these ligands transmit JAK-STAT signaling). **B**, Statistical analysis of GSC number during the dedifferentiation process in different (control vs. regeneration conditions) genetic backgrounds. In case of* upd1-RNAi, *\*: p<0.05, in case of* upd3-RNAi, *\*\*: p<0.001. ±S.D. indicates mean deviations*.

When an immune-depleted environment was induced, a TUNEL assay was applied to uncover increases levels of cell death, which occurred exclusively during the regeneration process (Fig. 6). Similar results were obtained when we inhibited the JAK-STAT signaling pathway. This Both *HS* and *UAS-rpr* lead to defects in the regeneration process. GSCs indicated by dotted suggests a link between the number of apoptotic SGs and JAK-STAT signaling. In the absence of hemocytes, there was no increase in apoptosis even under normal condition. Similar to that, there was no TUNEL-positive structure during the normal regeneration process (control). However, when hemocytes were eliminated, the number of TUNEL-positive cells increased significantly under regeneration-inducing conditions, indicating higher levels of apoptotic cell death. Similar results were obtained when *upd3* was specifically silenced in hemocytes. Together, these results suggest that hemocytes contribute to the regeneration process and their absence leads to the activation of caspase-independent apoptosis.

**Figure 6.**
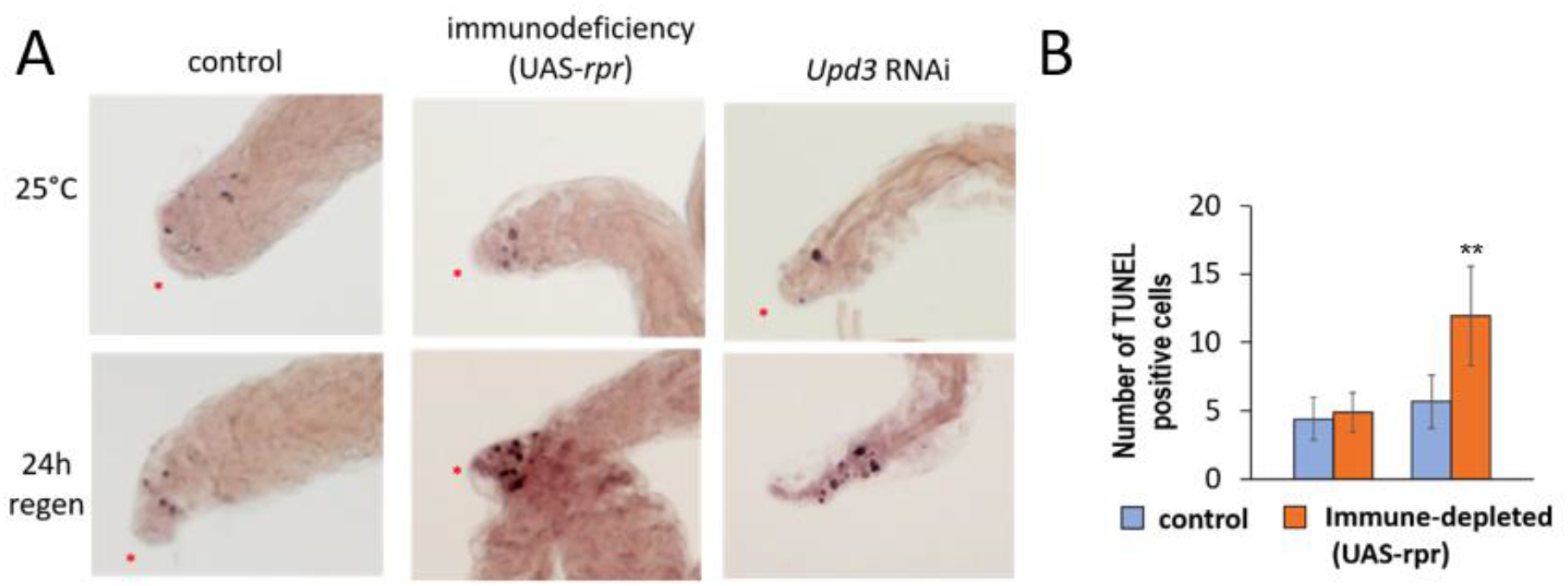
upd3 *downregulation in hemocytes increases the number of TUNEL-positive structures during the dedifferentiation process*. ***A**, TUNEL staining indicates an increase in the number of TUNEL-positive cells under regeneration-inducing conditions. This indicates an increase in apoptotic cell death. Red stars indicate the apical end of the testis. **B**, Changes in the number of TUNEL-positive structures under control (the first two columns) and regeneration-inducing (the second two columns) conditions, and in normal (blue) and immune-depleted (*UAS-rpr, *orange) states. During regeneration, a significant increase can be observed in the number of TUNEL-positive structures (**: p<0.01). ±S.D. indicates standard deviations*.

The multifunctional hemocytes participate in the removal of apoptotic corpses of SG origin, in which Omi/HtrA2 plays a key role^25^.

We could uncover a link between Omi and JAK-STAT signaling; s*tat92e* encodes a transcription factor for JAK-STAT signaling and its downregulation in the germline resulted in enhanced levels of *omi* expression (Fig. 7).

**Figure 7.**
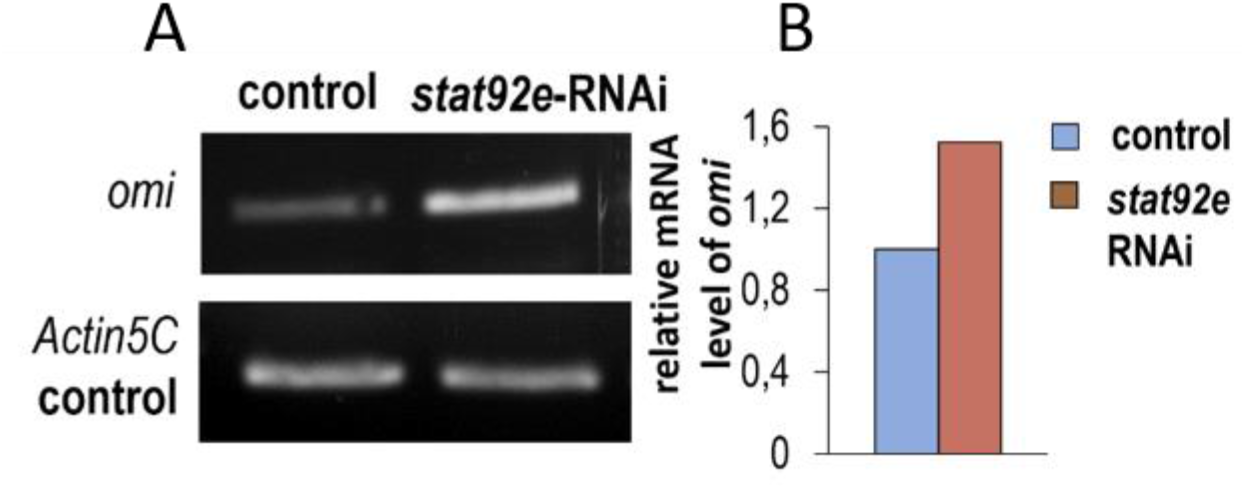
omi *expression in the germline is repressed by JAK-STAT signaling*. ***A**, Expression levels of omi are determined by RT-qPCR. Control vs. stat92e (serving as a transcription factor for JAK-STAT signaling) deficient backgrounds were compared*. stat92e *was downregulated in SGs.* omi *expression increases when* stat92e *is downregulated. **B**, Comparing relative transcript levels of* omi *in control (blue) vs.* stat92e *defective (brown) backgrounds.*

During GSC regeneration, *omi/HtrA2* expression was changed as revealed by a semi-qPCR analysis (Fig. 8). At the beginning of regeneration (0-2 h after induction), *omi* expression became lowered, while it was increased after 24 h of regeneration (Fig. 8).

**Figure 8.**
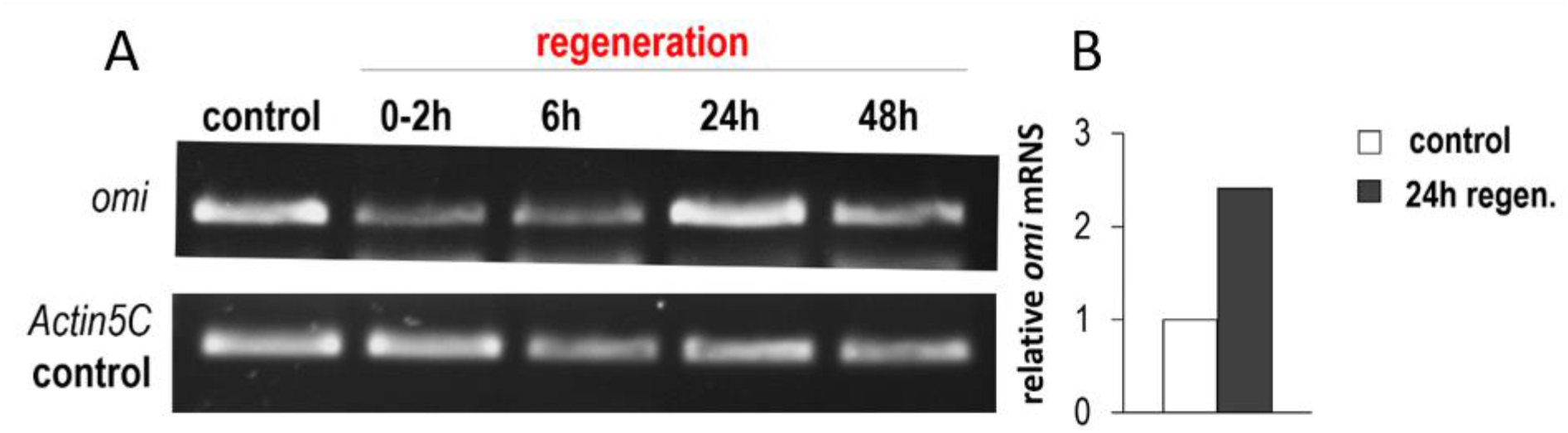
omi *expression changes at different stages of regeneration.* ***A**, Semi-qPCR analysis showing that* omi *expression decreases at early stages of regeneration, while becomes increased followed by 24 h of regeneration.* Actin5C *was as an internal control. **B**, Relative amounts of* omi *transcripts were determined by RT-qPCR. Control vs. 24 h regeneration were compared.*

Then, we examined the potential role of *omi* in GSC regeneration, by using fluorescent microscopy. We found that inhibiting omi transcription in the germline blocks the dedifferentiation process. Based on these results, Omi is required for the regeneration of GSCs (Fig. 9).

**Figure 9.**
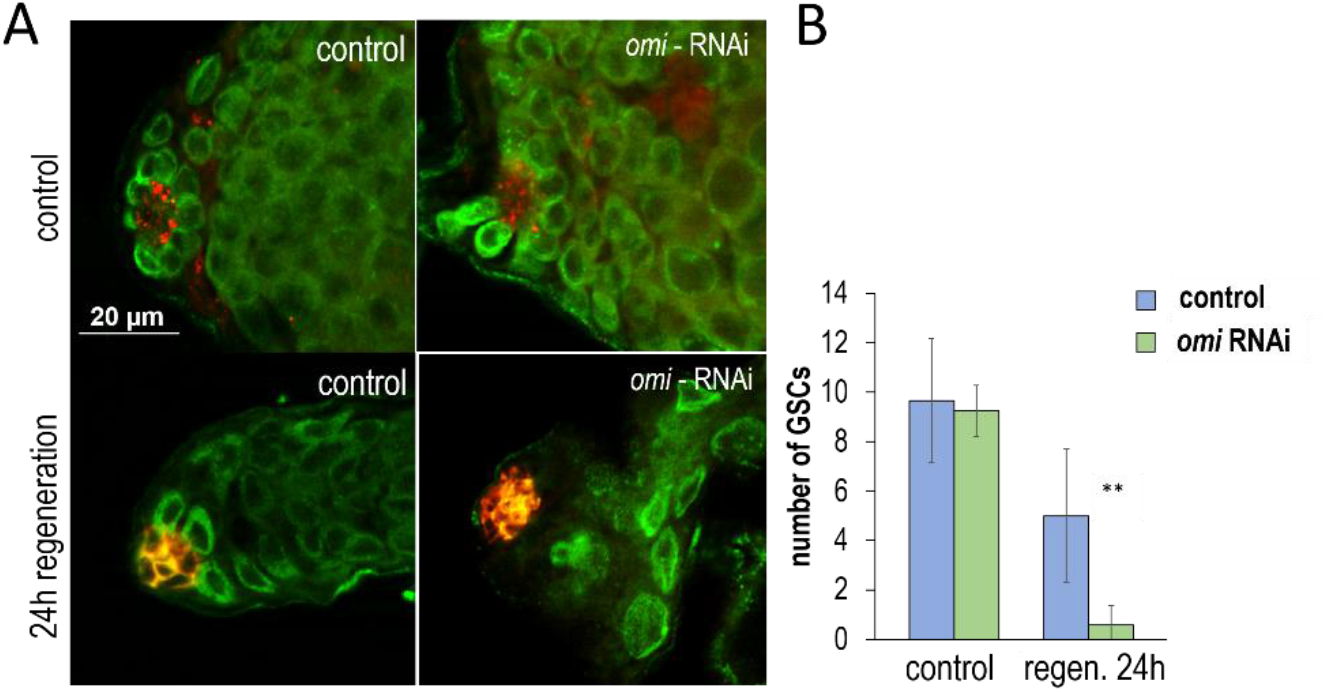
*Downregulation of* omi *in the germline interferes with GSC regeneration*. **A**, Dediifferentiation of SGs into GSCs in control vs. Omi defective backgrounds. Immunhistochemistry was applied: anti-Vasa indicates germline cells, while anti-FasciclinIII shows hub cells**. B**, Diagram showing the number of GSCs during the regeneration process. After 24 h, a significant level of difference was observed (**: p<0.005). Blue indicates control, green labels Omi-depleted states. bars indicate ±S.D.

Using a TUNEL assay, we tested changes in programmed cell death in Omi defective background. The results showed that *omi* downregulation leads to an increase in the number of cells undergoing apoptosis (Fig. 10). However, the increase was evident under both normal and regeneration-inducing conditions, but the latter was more significant as compared with the former condition.

**Figure 10.**
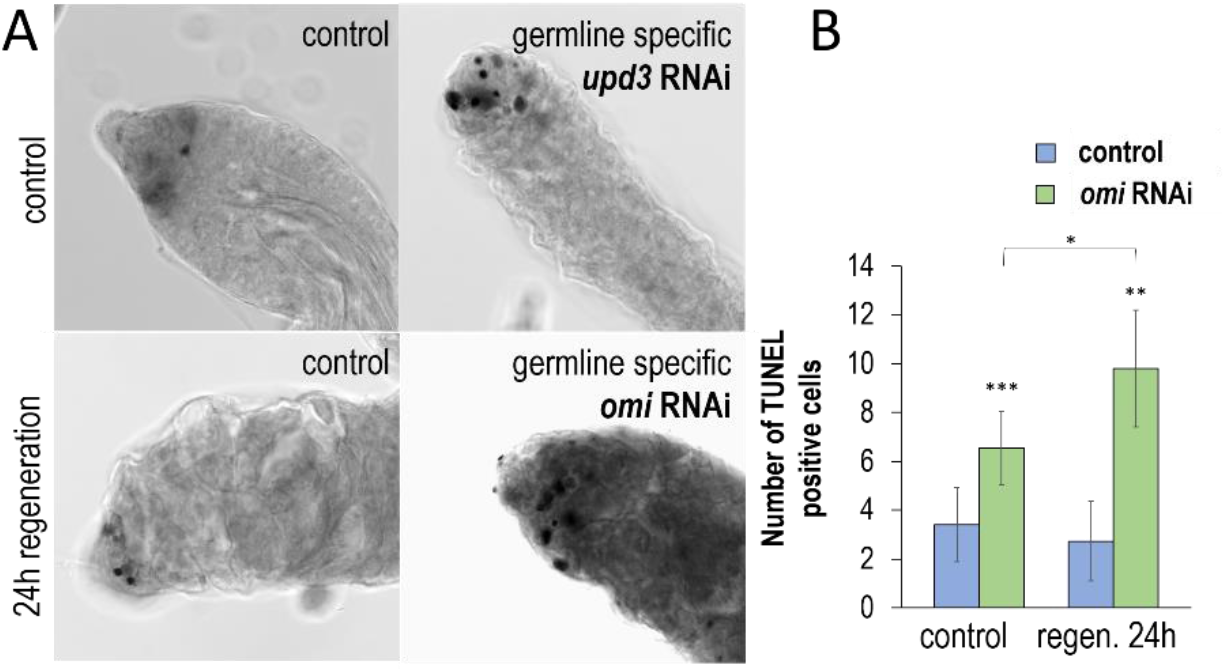
omi downregulation increases the number of TUNEL-positive structures during GSC regeneration. **A**, TUNEL assay showing that Omi depletion elevates the number of TUNEL-positive structures both under normal and regeneration-inducing conditions. **B**, However, the increase was more significant under the regeneration-inducing conditions. *: p<0.05; **: p<0.005; ***: p<0.001; bars indicate ±S.D.

During regeneration of GSCs, we performed a LysoTracker Red staining and could observed an increased level of acidic compartments, which indicates elevated autophagic activity (Fig. S2). These results may result from two processes. Either autophagy became hyperactivated or a later stage of the process was inhibited, causing the accumulation of early autophagic structures (phagophores and autophagosomes). To distinguish between the two alternatives, we quantify Ref(2)P levels, which orthologous to mammalian p62^26^. Ref(2)P/p62 serves as a substrate for autophagy, so its levels inversely correlate with autophagyic activity. We found that when *omi* is downregulated under normal conditions, Ref(2)2/P62 levels decrease significantly (Fig. 11). However, Omi deficiency significantly increased the number of Ref(2)P/p62-positive structures under regeneration-inducing conditions (Fig. 11). These results show that during GSC regeneration Omi is required for the activation of the autophagic process. In the *Drosophila* male germline, decreased levels of autophagy may lead to an increased apoptotic response (Figs. 10 and 11)^27^.

**Figure 11.**
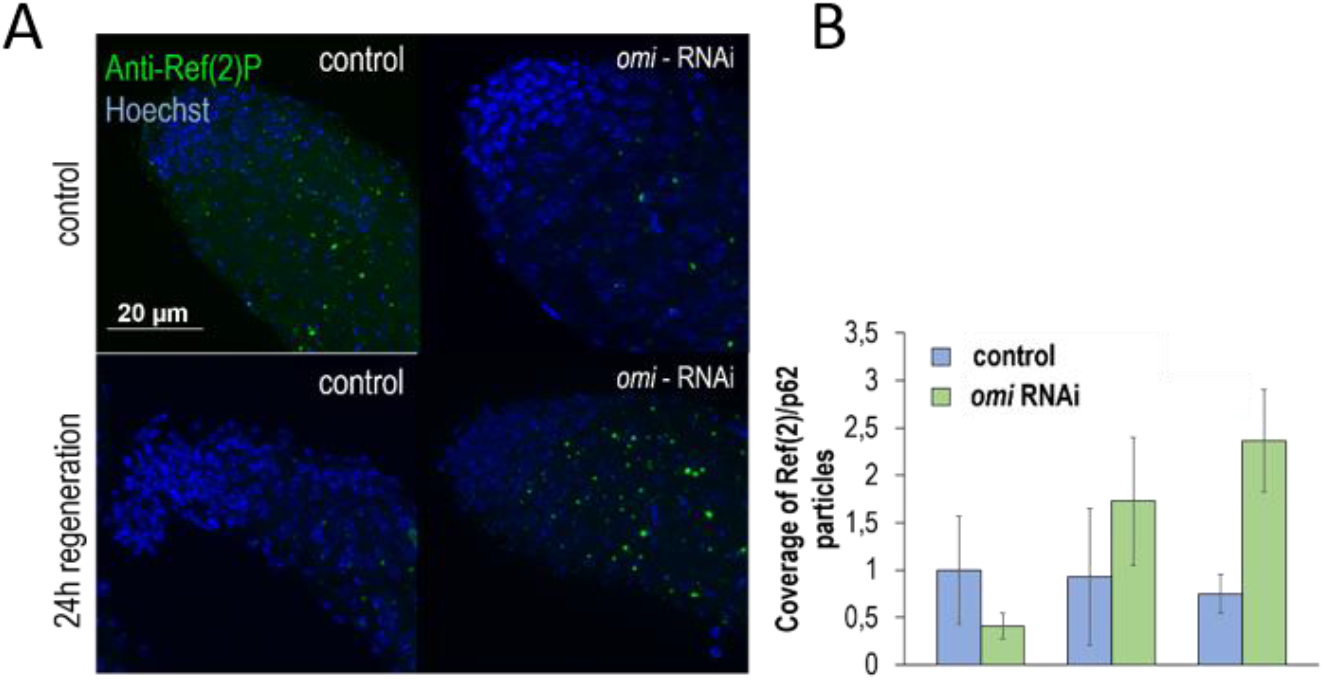
Silencing omi represses the process of autophagy. **A**, We applied anti-Ref(2)P (green) mark that is known as a substrate of autophagy, which made it possible to observe that silencing omi repressed autophagy. While under regeneration conditions Ref(2)P positive dots increased. **B**, Diagram showing the coverage of Ref(2)P positive particles under normal (blue) and regeneration (green) conditions. Bars indicate ±S.D.

## Conlusions

During the dedifferentiation process, hemocytes play multifunctional roles. First, these cells are capable of inducing signaling pathways required for the dedifferentiation of SGs into GSCs. Upd1 and Upd3 function as ligands for JAK-STAT signaling, and our results obtained here indicate that they are omitted by hemocytes. Thus, JAK-STAT signaling is activated in SGs by hemocytes. Upds can transmit through either Cyst cells (in which JAK-STAT signaling becomes activated to cause Upd expression toward SGs) or directly reach SGs. It is possible that at this time Cyst cells do not surround dedifferentiating germline cells.

During regeneration, JAK-STAT signaling may play a dual role in SGs. First, it may activate autophagy, which is required for cytoplasmic remodeling of SGs undergoing dedifferentiation. Second, the pathway may inhibit apoptosis through repressing a mitochondrial serine protease, HrtA2. In the presence of JAK-STAT signaling Omi expression becomes lowered, and its low levels are required for regeneration. Omi may play an important role in maintaining the balance between autophagy and programmed cell death (Fig. 12). Together, results obtained by this study may contribute to a better understanding of mechanisms underlying tissue regeneration, and its regulation by immune cells.

**Figure 12.**
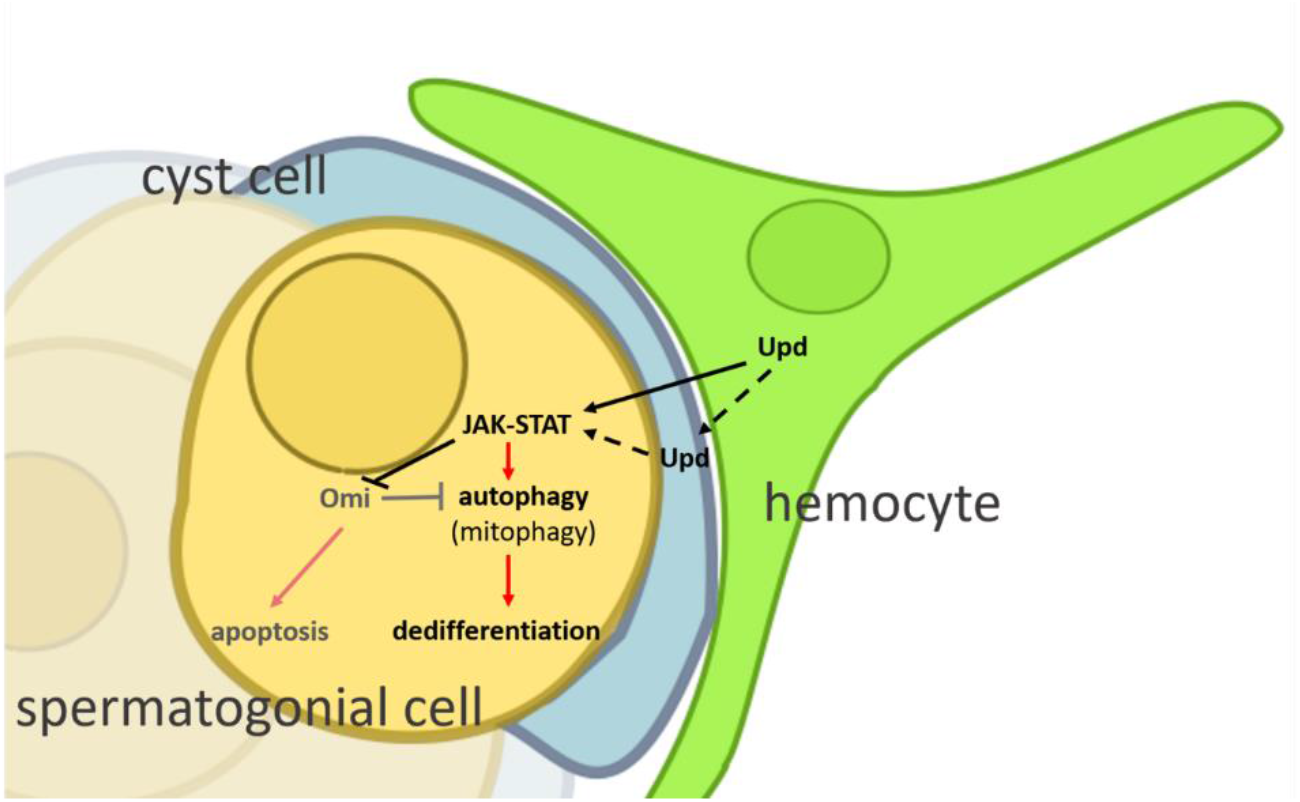
Multiple functions of immune cells in tissue regeneration. *Hemocytes are able to transmit ligands of signaling pathways to germline cells. Such a pathway is represented by JAK-STAT signaling, which may have a dual role in the* Drosophila *male testis. First, it maintains the number of GSCs. On the other hand, it induces dedifferentiation of SGs into GSCs. The signaling pathway also acts as a relay of whether dedifferentiation occurs through autophagy, or caspase-independent apoptosis occurs through Omi*.

## Materials and Methods

### *Drosophila* strains and their maintenance

For the maintenance of the used frutifly strains the compliants and characterications of the food were published in the article below.^28^

Strains that were provided by Bloomington Drosophila Stock Center-ből (BDSC): *w[1118]* (#5905), *P{w[+mC]=hs-bam.O}18d, w[1118]* (#24636)*, y[1] w[67c23]; P{y[+mDint2] w[BR.E.BR]=SUPor-P}crq[KG01679]/CyO* (#14900), *w[*];y[1] sc[*] v[1] sev[21]; P{y[+t7.7] v[+t1.8]=TRiP.HMC03843}attP40* (#55165), *y[1] sc[*] v[1] sev[21]; P{y[+t7.7] v[+t1.8]=TRiP.HMS01328}attP2* (#34340), *w[1118]; P{w[+mC]=HmlGAL4.Delta}3* (#30141), *y[1] sc[*] v[1] sev[21]; P{y[+t7.7] v[+t1.8]=TRiP.HMS01244}attP2* (#34899), *y[1] v[1]; P{y[+t7.7] v[+t1.8]=TRiP.HMS00035}attP2* (#33637), *y[1] sc[*] v[1] sev[21]; P{y[+t7.7] v[+t1.8]=TRiP.HMS00545}attP2* (#33680), *w[1118]; P{w[+mC]=Hml-GAL4.Delta}3* (#30141*), y[1] v[1]; P{y[+t7.7] v[+t1.8]=TRiP.HMC03230}attP40* (#51483), y[1] v[1]; P{y[+t7.7] v[+t1.8]=TRiP.HMC03230}attP40 (#51483) *amosTft/CyO; BamGal4* was provided by Sinka Rita (University of Szeged, Department of Genetics), All other lines were established for this study

### Protocol of Dedifferentiation

We performed experiments on the *Drosophila* testis in order to observe regeneration via dedifferentiation because GSC regeneration can be activated under laboratory conditions. Based on the *hs-bam, bamGal4 expression system*, *bam* is driven by a heat inducible promoter (Hsp70) ectopically expresses the main germline regeneration factor Bam (bag of marbles)^17^. Under normal conditions, *bam* is silenced in the germ line. Due to ectopic expression of *bam*, it is expressed in GSCs and makes differentiation possible. By finishing ectopic expression of Bam, dedifferentiation of SGs (and GBs) into GSCs can be activated. This process is termed as the regeneration of early spermatogenesis. After the termination of ectopic expression recovering GSCs is possible by only spermatogonial dedifferentiation^7^.

Regeneration protocol step by step:

Fruitflies were maintained at 25°C unless they were used to the regeneration protocol
Day 1: 3×30 min heat shock by applying 37°C water bath. In between the water baths at least 2 hours needs to be elapsed and maintained at 29°C
Day 2: Besides the water baths fruitflies are maintained at 32°C
Day3: The fruitflies are taken back to the 29°C termostate.

Applied control groups:

Fruitflies were maintained at 25°C temperature in a termostate.
2 regeneration groups were used according to how many hours are passed after terminating the ectopic expression of *bam*:
  1. 24h: After 3 days of heatshock then keeping them at 25°C temperature for 24 hours
  2. 6h: Same as 24h but keeping them at 25°C temperature for 6 hours
  3. *bam* overexpression (HS): maintaining this group at 29°C for 3 days. Maintaining a high level of bam expression, thus preventing regeneration.

### Immunohistochemistry

We used the following staining protocol for the labeling of germline cells and GSCs.^29^

First antibodies: anti-VASA: DSHB, (mouse, 1: 50), anti-FasciclinIII: DSHB, (rat,: 1: 25)

Secondary antibody: anti-rat: Life Technologies #A21210, Alexa Fluor 488, (rabbit, 1:500), anti-mouse: Life Technologies #T862, Texas Red, (goat, 1:500).

### Fluorescent microscopy

A fluoreszcens képek készítése: Images were captured with Zeiss Axioimager Z2 upright epifluorescent microscope equipped with ApoTome2 semi-confocal upright. We used the following softcovers: ZEN 3.1

Images were captured with Zeiss Axioimager Z2 upright epifluorescent microscope equipped with ApoTome2 semi-confocal upright. We used the following._programs for evaluation of images: ZEN 3.1 (blue edition) and Fiji ImageJ 1.8.0_77.

### Electron Microscopy

Electron microscopy images were taken with JEOL (1011; JELOL, Tokyo Japan) transmission electron microscope equipped with Olypmos Morada digital camera, and we used iTEM software ^30^

We used the following solution during fixation:

2,4 ml 16% formaldehyde (final concentration: 3,2)
480 μl 25% glutaraldehyde (1%)
0,12 g sucrose
0,00435 g CaCl2
Na-cacodylate 0,1M 7,4pH

### qPCR reaction and applied primeres

#### 1. RNA isolation and cDNA synthesis

The whole amount of RNA was extracted from 20 testes using TRI reagent (Sigma, T9424), then purifying RNA was conducted by Directzol RNA MiniPrep kit-tel (Zymo Research, R2050) and was performed as described in manufacturer’s protocol. The single strand cDNA synthesis was provided by RevertAid First Strand cDNA Synthesis Kit (Thermo Scientific, K1622).

#### 2. Real-time quantitative PCR

o Using LightCycler 96 System (Roche, FastStart Essential DNA Green Master, 06402712001) under the circumstances of denaturation: 10 minutes at 95°C; then 45 amplification cycles (10 sec,95°C; 10 sec, 58°C and 20 sec, 72°C). Primers used for the amplification:

- As controll: *Actin5C*: forward – 5’-GGA TAC TCC CGA CAC - 3’; reverse – 5’- GAG CAG CAA CTT CGT CA - 3’
- *chinmo*: forward – 5’ - AGC AGT TCT GCC TCA AAT GG 3’; reverse – 5’- AGA TCG GCG AAC TTC TTT GA - 3’
- *socs36e*: forward – 5’ - TCG TCG AGT ATT GCG AAG TG - 3’; reverse 5’- CTG CTC CCA TTG AAA GTG CT - 3’
- *Omi/HtrA2:* forward – 5’ – CTT TGC GCA TAC AGG TGA AC – 3’; reverse – 5’CGC TGC GTT GAA CTG ATT AC – 3’
- *mRpL11:* forward – 5’ – CAC CAA CAC ATT TCG CTT TG – 3’; reverse – 5’ – TTC CCG CAG GTA TAT TCG TC – 3’

PCR products were detected by a fluorescent, DNA-binding (double strand) SYBR Green dye (Roche FastStart Essential DNA Green Master 11560300).

Verifying the specificity of the reaction of qPCR was provided by analyzing the melting curves. By normalizing the values of the PCR threshold cycles, it was possible to identify the average of relative mRNA level.

### TUNEL – assay

After the pre-fixation protocol, we followed the manufacturer’s instructions.^31^ We used the following materials during TUNEL-assay: ApopTag kit(Chemicon/Millpore, Billerica, MA, USA, S7105, S7106, S7107).

Testicles were fixed in 4% formaldehyde (diluted with PBS) for 20 minutes and washed 3 times in cold methanol. We incubated samples in −20°C cold methanol for 24 hours.

### Statistics

We used the RStudio (Version 1.2.5033) program for statistical analysis..

Lilliefors test were used to know that the distribution of samples examined is normal or not. If it was normal, F test was performed to compare two variances. If the variances were equal, two-sample Student’s t-test was used otherwise t-test for unequal variances was applied. If the distribution of a sample is not normal, Mann-Whitney U test was performed

## Supporting information

Supplemental image 1 and 2

## Acknowledgments

We thank Miklós Erdélyi and Rita Sinka for them advise, and thank Gábor Juhász for the Drosophila stocks.

## Funding

Supported by the ÚNKP-19-3 New National Excellence Program of the Ministry for Innovation and Technology.

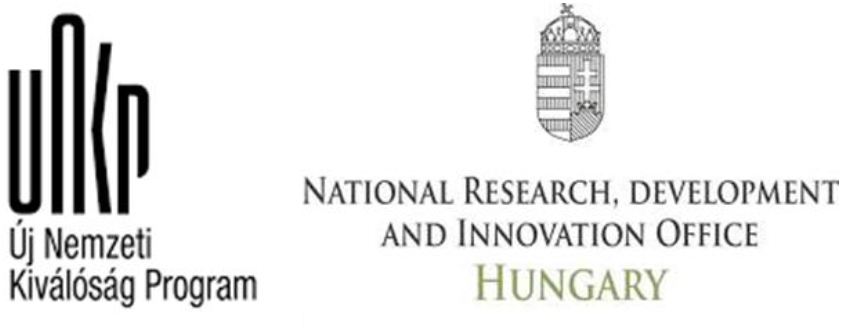

