## Supplemental image 1 and 2 for "Role of hemocytes in the regeneration of germline stem cells in *Drosophila*": supplementary.pdf

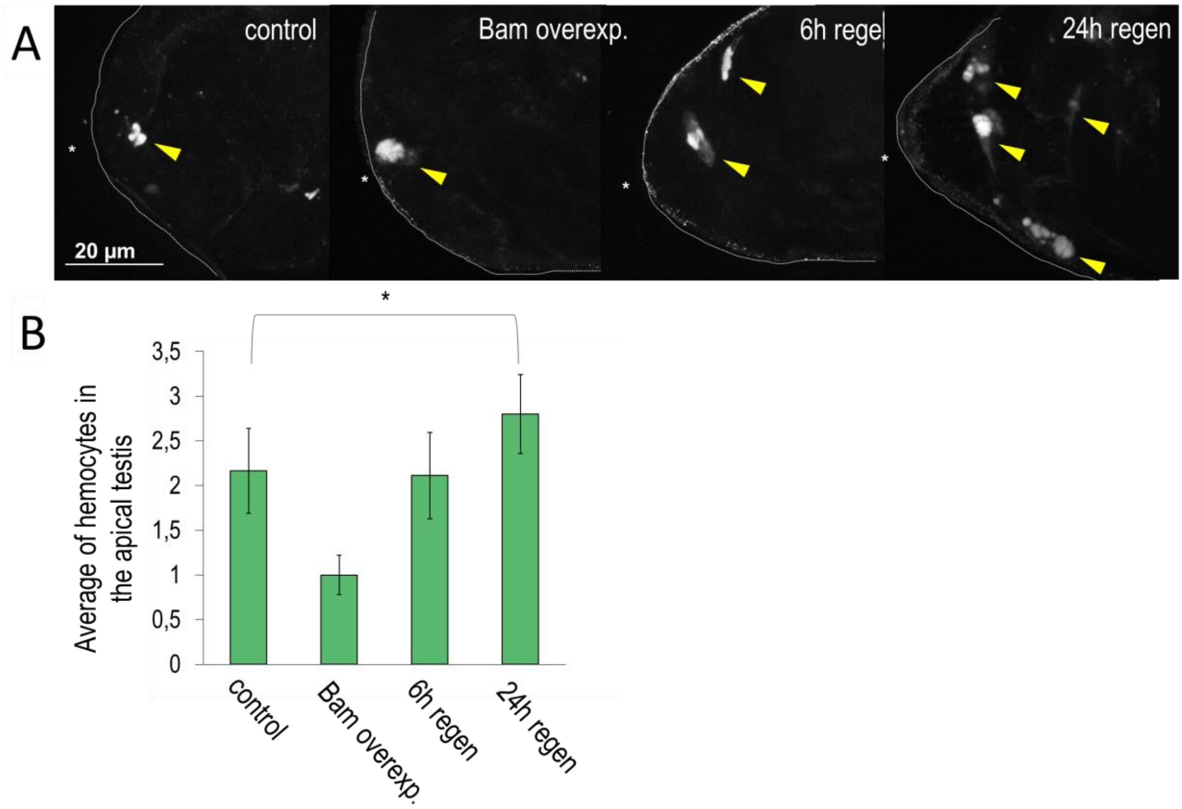


***Supplementary figure 1. Number of hemocytes is increasing during regeneration. A**, Applying Hml-GFP transgenes (UAS-myr-GFP is driven by Hml-Gal4), we observed that hemocytes are recruited to the site of GSC regeneration. The yellow arrowhead indicates the hemocytes. **B**, Diagram showing the coverage of LysoTracker Red positive particles under normal, bam overexpressed and regeneration (6h and 24h after the begin of regeneration) conditions. Bars indicate ±S.D.*

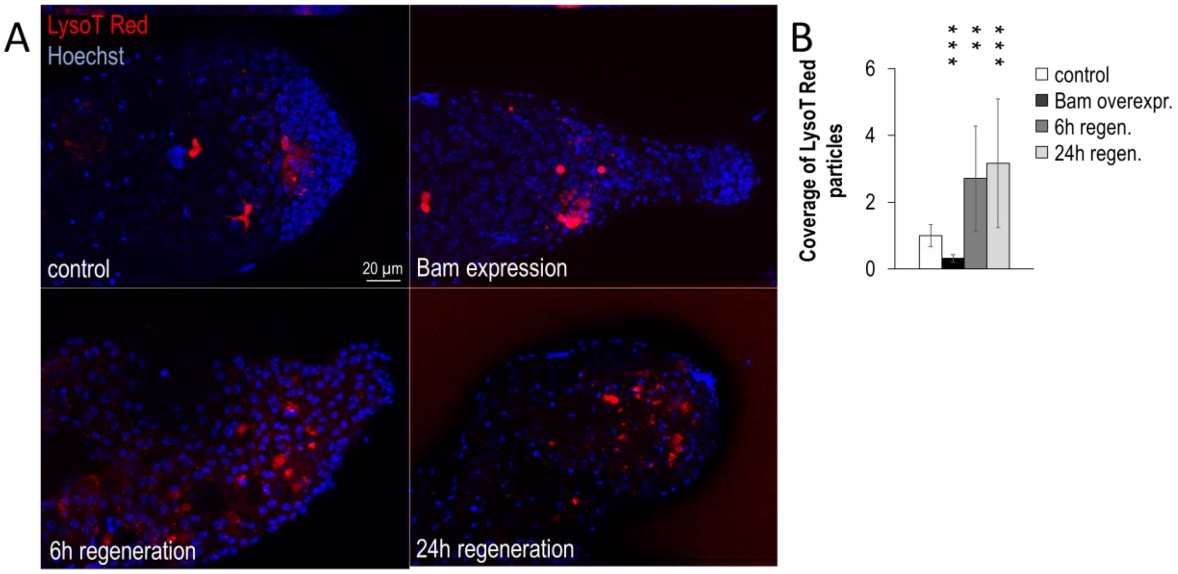


***Supplementary figure 2. During regeneration, the abundance of acidic compartments is increased. A**, We found an increased number of acidic compartments (LysoTracker Red - red) during regeneration. The blue color indicates the nuclei (Hoechst staining). **B**, Diagram showing the coverage of LysoTracker Red positive particles under normal, bam overexpressed and regeneration conditions. Bars indicate ±S.D.*
